# A Constant Proportion in Protein Structure

**DOI:** 10.1101/500025

**Authors:** Francisco Javier Lobo-Cabrera

## Abstract

The principles governing protein structure are largely unknown. Here, a structural proportion universal (R^2^ = 0.978) among proteins is reported. The model variance is shown to be independent from protein size, secondary structure composition, compactness or relative surface area. The structural characteristic under study --named here QUILLO-- quantifies residue-type spatial clustering. In this way, polar, hydrophobic, acidic and basic residues are evaluated individually and their values added up. For the analysis, all X-Ray currently determined structures deposited in the Protein Data Bank were studied. The QUILLO proportion offers for the first time an *a priori* protein prediction quality-check. Indeed, predictions with unexpected proportion values correspond to low ranks in the CASP12 experiment. The reason behind a specific, constant rule for protein folding remains unknown.

## Introduction

Biomolecules, such as nucleic acids and proteins, exert their biological function in a 3D environment [1]. Therefore, comprehension of their spatial characteristics constitutes an active area of research [2]. For proteins, this is relevant as they carry out most cellular functions, such as structural functions, catalysis of biochemical reactions, transport or signaling [3] [4] [5].

The first protein structure determined was that of myoglobin in 1958 [6]. Since then, over 125,000 protein structures have been determined [7][8]; mostly by X-Ray crystallography (~90%) and Nuclear Magnetic Resonance (~8%) [7][8]. Simultaneously, a variety of simulation techniques have been developed that allow prediction of tertiary conformation based on primary sequence [9] [10]. Additionally Molecular Dynamics and Quantum Mechanics methods enable refinements of the obtained structures [11] [12].

Despite the advancements in the field, prediction of three-dimensional structure remains challenging [13]. For example, it is known that proteins with similar primary sequence may have different 3D arrangements [14]. Also, some proteins change their conformation when interacting with specific substrates [15] or their structure can be modified by interaction with solvent molecules [16]. This emphasizes the need for more knowledge on protein structure and the generation of more sophisticated prediction methods. In turn, this new knowledge on structure will benefit understanding of protein functioning.

One way to make new discoveries on protein structure comprises statistical analysis of already available data. Currently, there are multiple databases containing polypeptide information. Examples include the Protein Data Bank [7][8] –with experimental structural data, the DSSP database [17][18] –containing protein secondary structure-- or BRENDA [19][20] – an enzyme database. In these sources, the user can simply enter a specific query (e.g protein name or id) and receive the information related to that query as output. The referred databases also allow systematic access by users, so that an automatic protocol can retrieve information for multiple queries. In this fashion, they enable direct analysis of a large number of polypeptides. The power of the aforementioned databases, combined with proper analysis tools, can render new insights into proteins. As an illustration, both CATH [21] and SCOP2 [22] have used data from the Protein Data Bank to establish structural and evolutionary relationships between protein domains.

Databases may also be employed to extract potentially constant features of proteins. Up to know, only a few characteristics have been shown to be general among polypeptides. The 20 standard amino acids, the nature of the peptide bond [23], a limited number of secondary structure conformations [24] or the organization of proteins in domains are among them [25] [26]. These elements have helped gain understanding of general polypeptide structure and functioning. However, not all of them are truly universal. For example, over 120 additional amino acids have been identified as naturally occurring in proteins [27], some proteins do not contain domains [26] and the secondary structure relative composition is highly variable from protein to protein; where some proteins may contain only alpha helices and not beta sheets or vice versa [28]. Noteworthy, these general features have been identified by traditional means and not by statistical screening of databases. Hence, the possibility exists that there are hidden universal traits that only database mining can unveil.

In order to search in databases for unraveled constant characteristics it is necessary first to define those characteristics. In this case, the number of possible designs is enormous; one can measure practically any kind of physical, chemical or biological protein parameters, or even proportions or ratios between them. For example, number of residues, compactness, electric charge, aliphatic index… Again, the existing databases provide the necessary input information. There are already tools for the computation of multiple of these parameters, such as ProtParam [29], PDBParam [30] or Vossolvox [31]. Once the characteristic to measure is defined, the polypeptides can then be scored. If the score is the same for all proteins, in principle the characteristic or trait is shown to be constant. Of course, since proteins are a highly diverse group of molecules [32] it is expected that only a few characteristics will be invariant.

In this article, a chemo-spatial parameter for polypeptides is presented. Referred as QUILLO (**Qu**antitative **I**nternal **L**ow **L**evel **O**rder) it quantifies the presence of spatial clusters of the same type of amino acid. Once defined, an analysis was conducted that included all X-Ray determined entries in the Protein Data Bank. Remarkably, the QUILLO value for a protein structure is highly predictable. In fact, it is only required to take into account the number of amino acids it contains. This demonstrates the existence of a QUILLO-related proportion, universal apparently among proteic structures.

Albeit that the QUILLO proportion is shown to be highly constant, there is still some limited variation among proteins. This variation could be simply stochastic, but several analyses were performed to try and correlate those differences with key structural characteristics. In this manner, the proportion was evaluated in its relation with secondary structure composition, protein compactness and surface area per residue. The aim was to test the independence between the QUILLO proportion and each characteristic.

## Results

### The QUILLO proportion

The first step in this work involved calculating QUILLO (see Methods) in a large set of proteins. For this purpose, the Protein Data Bank (PDB) archive --specifically those entries determined by X-Ray diffraction-- was employed. Currently, PDB includes structural data of more than 130,000 polypeptides and peptides. Of these, approximately 90% have been resolved by X-Ray diffraction [7][8], a technique that offers very high atomic resolution [33]. Results of the analysis of X-Ray entries from PDB show that a structural proportion exists, uniform (R^2^ = 0.978) among proteins. This proportion relates the QUILLO value and the number of residues of a protein. Specifically, it shows that QUILLO increases its value with the number of residues following a specific pattern (Figure 1).

**Figure 1.**
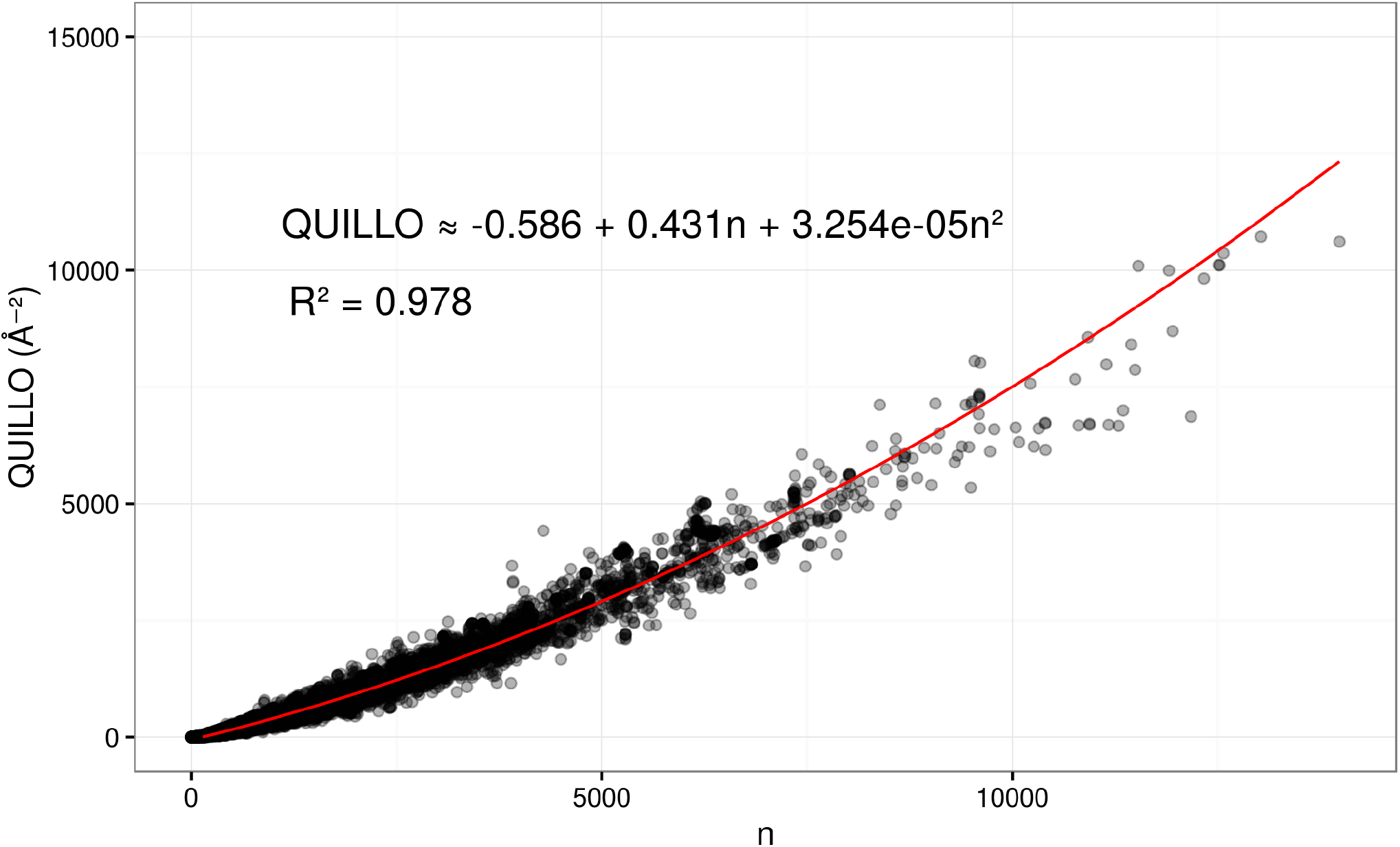
The QUILLO proportion. A simple linear model establishes the relationship between QUILLO and the number of residues (n). For the calculation, distances between residue space points are measured in Ångströms. A total of 111 out of 113081 entries (< 0.1% of the data set) were discarded as outliers.

The other defined parameter –QUILLO2-- was likewise assessed to search for a proportion. In this case, QUILLO2 showed lower correlation (R^2^ = 0.891) than QUILLO to the number of residues.

### Independence from different factors

In all the following cases, deviations from expected values (given the QUILLO proportion) are plotted against a certain protein characteristic. Deviations are simply the difference between the obtained QUILLO value and that predicted by the proportion. The input data set corresponded to the X-Ray entries of PDB. Specifically, secondary structure composition, protein compactness and surface area per residue were studied.

#### a) Secondary structure composition

Proteins display a limited number of secondary structure types, being alpha helix and beta strand the most abundant [34] [35]. To determine whether secondary structure affects QUILLO values, the percentage of residues in beta strand of each protein was calculated. Approximately, this percentage becomes inversely proportional to that of residues in alpha helix.

Figure 2A shows absence of correlation between secondary structure and the QUILLO proportion.

**Figure 2A.**
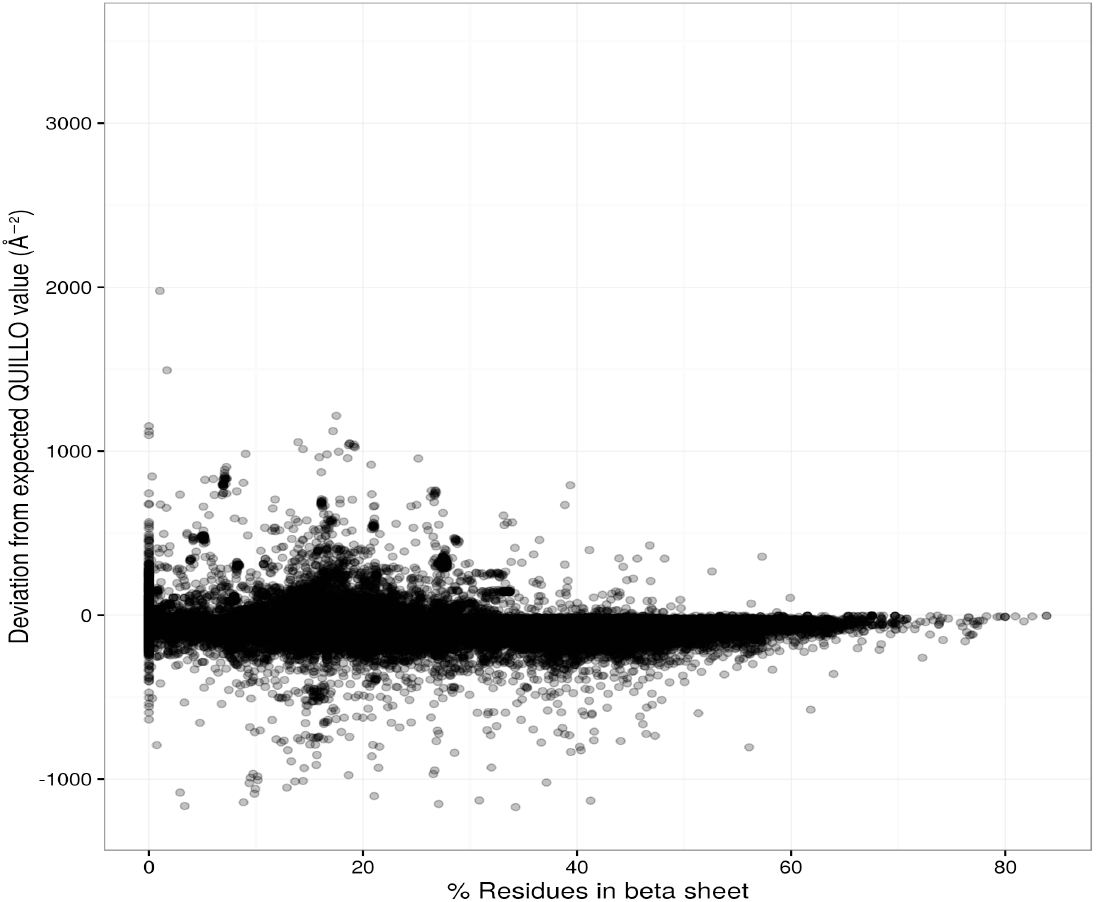
Independence from secondary structure. Plotting of the percentage of residues in beta sheet and the deviation from the expected QUILLO value.

#### b) Compactness

Protein compactness refers to the arrangement, more or less extended, of the residues in a protein. As QUILLO measures distances between residues, it was hypothesized that it may be dependent on the overall compactness.

As depicted in Figure 2B, there is very low correlation between the two variables, indicating little effect of protein compactness.

**Figure 2B.**
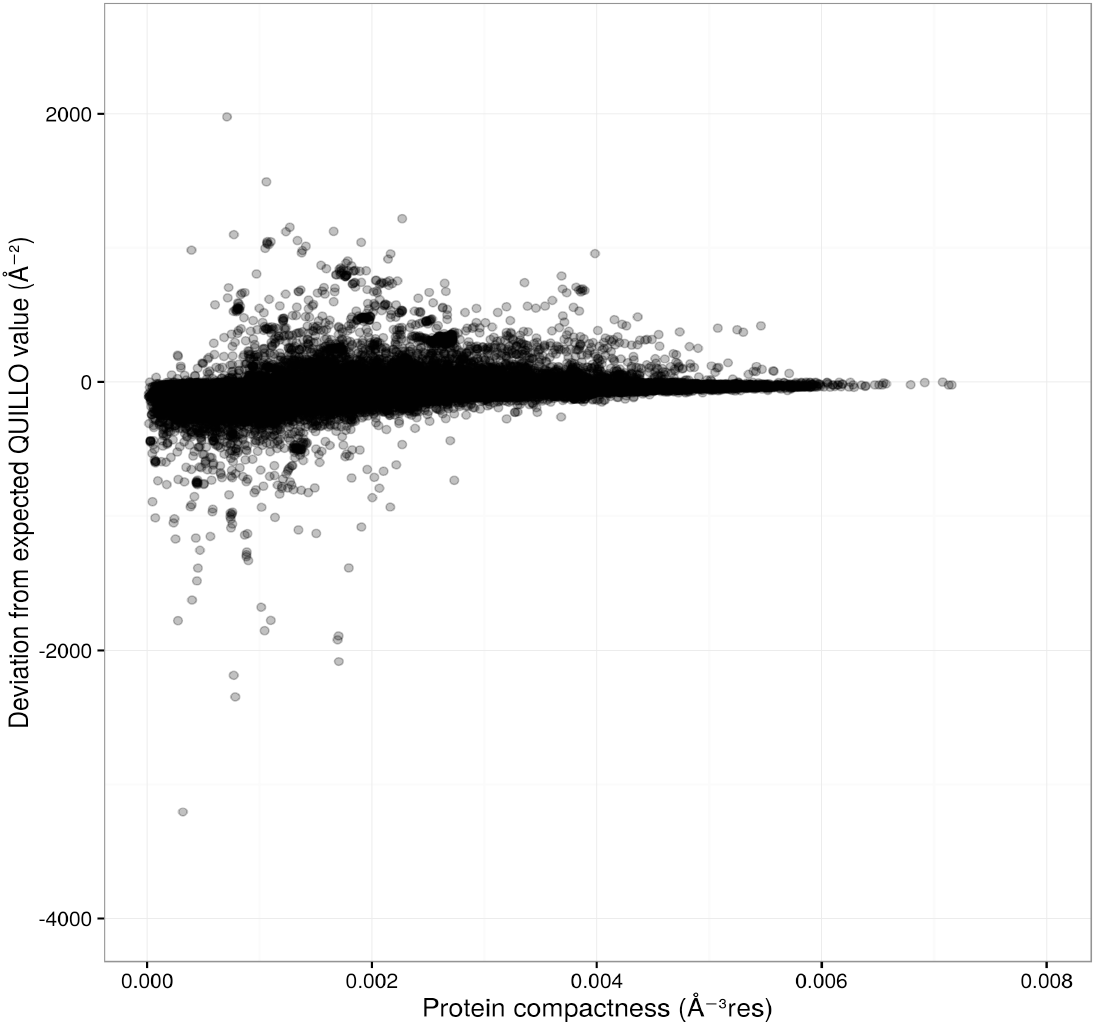
Independence from protein compactness. Protein compactness here is referred as the residue density in an imaginary sphere of minimum volume containing all the protein atoms. Note: One point whose compactness is > 0.06 Å^−3^ res was not plotted.

##### c) Surface area per residue

Another studied structural parameter was the surface area per residue. Polypeptides with higher proportion of buried residues; that is, residues not accessible by the solvent, will have lower surface area per residue.

As shown in Figure 2C, the spatial rule under study in this article does not depend on the surface area per residue.

**Figure 2C.**
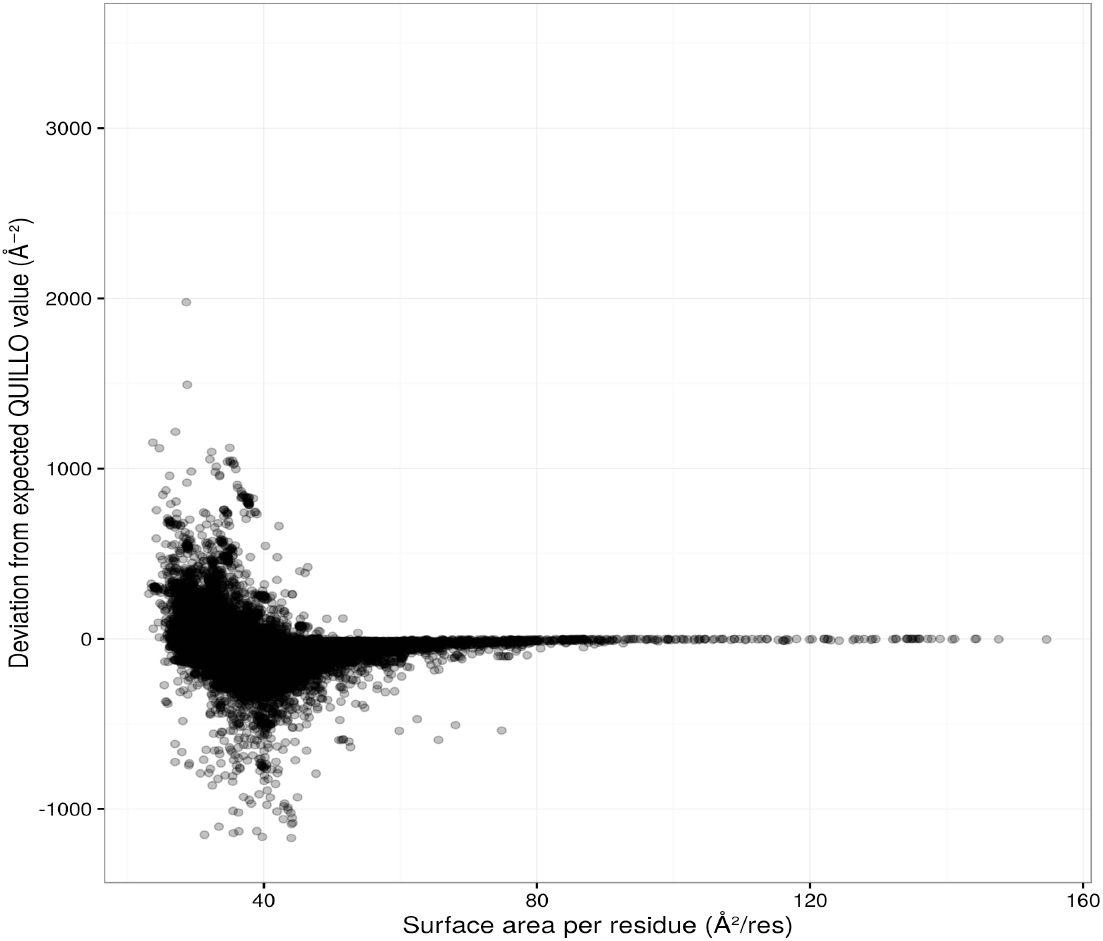
Independence from surface area per residue. The surface area for each protein was calculated with the 3V software [40], using a probe radius of 1.5 Å.

#### An *a priori* method for quality-check of structure prediction

A constant proportion enables in principle inference on structure. Therefore, an *a priori* method can be constructed to check the quality of structure predictions, without the need to compare with experimental results. To our knowledge, this is the first time a test of this kind is proposed.

To explore the use of the QUILLO rule as a quality-check, data from the CASP12 experiment [36] was analyzed. CASP12 [36] constituted a recent evaluation of computational methods for protein structure prediction. It ranked simulations from different research groups in relation to an obtained experimental structure. CASP12 assessed in this way predictions for a total of 53 protein targets. The goal here was to see whether there was a correlation between CASP12 rankings and deviations from expected QUILLO values.

Results show that i) the QUILLO proportion does not allow to compare between good predictions, but rather to ii) identify poor predictions. This can be explained by the fact that even the experimental structures do not have a QUILLO difference equal to 0. So, predictions of those structures do not necessarily need to have QUILLO differences ≈ 0 to be optimum. On the other hand, in the PDB X-Ray data set 90% of the cases have absolute difference < 111.6866 Å^2^ from the expected QUILLO value. In this manner, it would be relatively safe to discard (90% probability) those predictions displaying differences larger than 111.687 Å^-2^. This opens the door for *a priori* quality assessment of structure predictions. Accordingly, Figure 3 depicts how in CASP12 predictions surpassing the referred limit correspond to low ranks.

**Figure 3.**
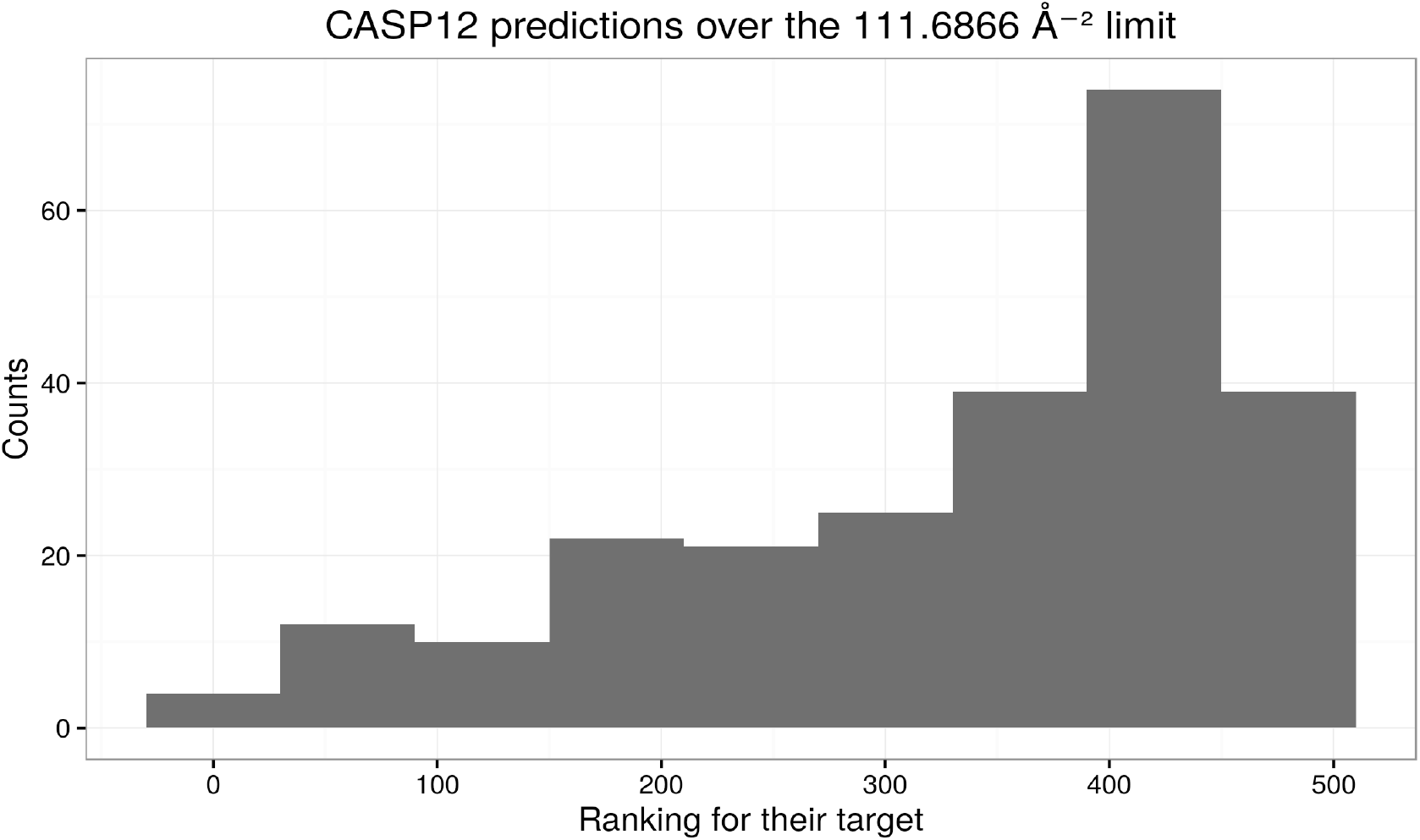
Checking the validity of the test on the CASP12 data. This diagram represents the rankings of those predictions surpassing the 113.2107 Å2 limit. Instead of a uniform distribution, the diagram shows that low rankings are favoured.

#### Origins of the QUILLO proportion

Here some hypothesis are for the factors creating the proportion. These include a thermodynamic and a biological perspective.

##### a) Thermodynamic approach

a general principle for protein folding strongly suggests free energy stabilization as a key factor. In order to test this theory, a thermodynamic variable was chosen for the variety of proteins in the PDB X-Ray data set. In this case, it was the temperature of the system, meaning the temperature at which the protein is found. A thermodynamic basis for the proportion could mean that system temperature is related with deviations from proportion expected values.

Figure 4 shows that no real correlation is found between system temperature and the QUILLO proportion. Nevertheless, this does not exclude thermodynamics as a key factor.

**Figure 4.**
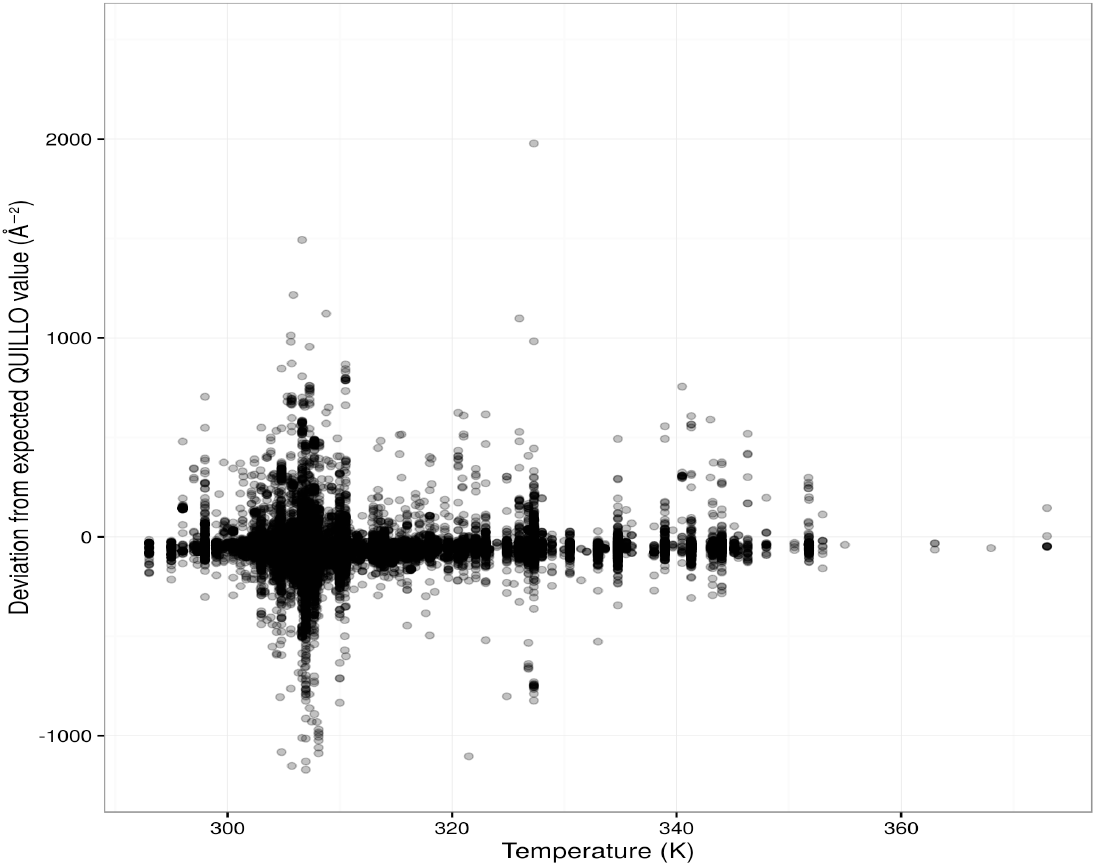
System temperature and the QUILLO proportion. System temperature corresponds to the approximate temperature at which the polypeptide is present in nature. It is calculated as the average optimum temperature of the organism’s enzymes included in the BRENDA database.

##### b) Biological approach

Other explanations regarding this rule may be biological. Nonetheless, they would have to be shared among proteins of all kind, as so does the rule.

According to its definition, high values of QUILLO imply that the protein contains large islands of amino acids of one type. This potentially can lead to strong interactions with molecules of the same chemical nature as those amino acids. Alternatively, low values of QUILLO would imply soft interaction with other molecules, as no great areas of amino acids of any type would exist. Life depends on molecular recognition, i.e interaction between molecules. These interactions must not be either too strong (molecules would aggregate unspecifically) or too weak (recognition would never occur). A common, shared rule like QUILLO could be the *radio frequency* at which molecular recognition among proteins --or among proteins and other molecules-- must happen.

#### Relation to cell adhesion and enzymatic activity

At the same time, deviations from QUILLO expected values exist for different proteins of same number of residues. This could be related to fine-tuning molecular interactions, and thus protein function. To check this, a GO-terms based analysis was conducted (see Methods).

The results show that only one GO-term (p-value < 0.05) is underrepresented in proteins with unexpected low QUILLO values and overrepresented in proteins with unexpected high QUILLO values. The referred GO-term is GO:0003824, which corresponds to catalytic activity. This is consistent with the hypothesis of increasing QUILLO values implying stronger molecular recognition.

By contrast, GO:0007155 is another GO-term related to molecular interactions, in particular to adhesins, that is not associated with either high or low QUILLO values.

## Methods

Easy-to-use Python code for all the calculations is available at <link>. On the other hand, ggplot2 [37] was employed in the generation of the graphics.

### QUILLO parameter definition

Given a polypeptide, let’s associate each of its residues with two characteristics; i) a position in space and ii) a chemical nature --acidic, basic, polar or hydrophobic. The position in space is established not as a volume but rather as a single point in space. This point is calculated as the geometrical center of the smallest cubic box encompassing the center of every atom in the residue (Figure 5B).

**Figure 5.**
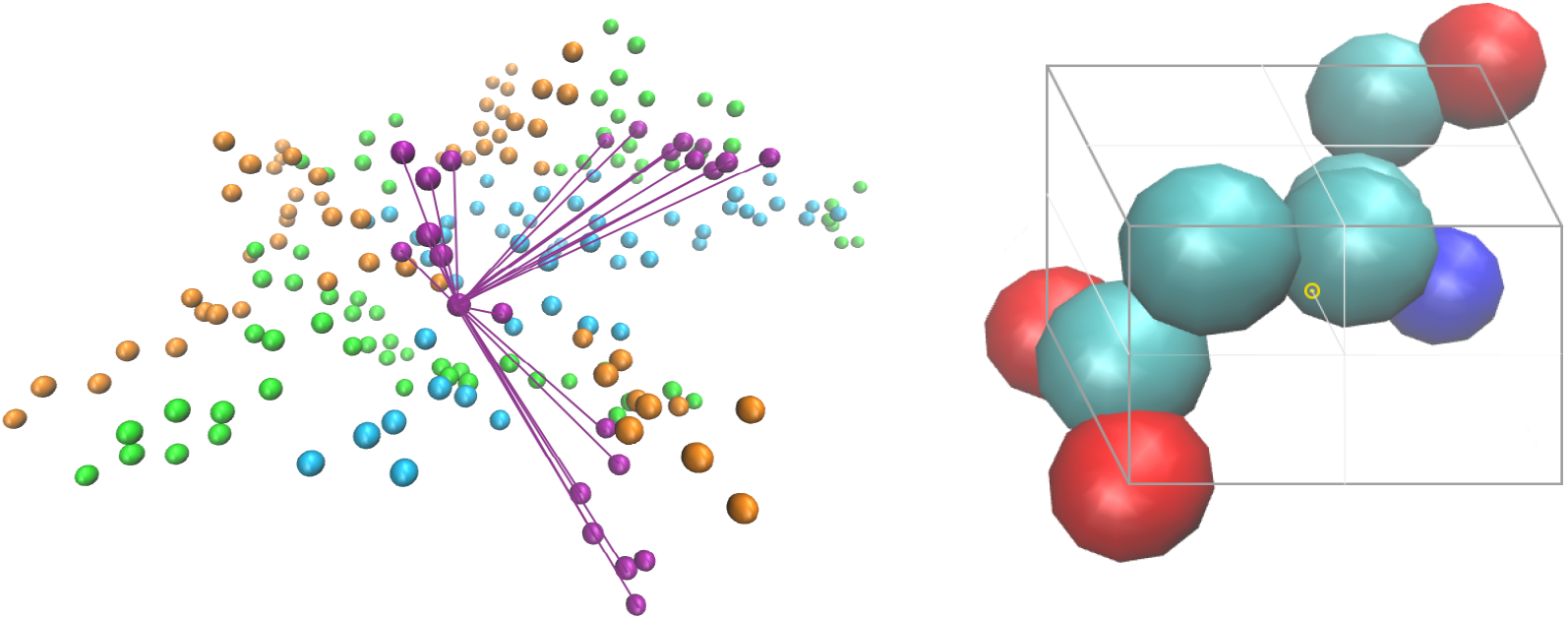
**A: Representation of the Euclidean distances of one residue with other residues of the same type.** Every sphere depicts the space point of a residue in an example protein, and is coloured according to its type (acidic, basic, polar or hydrophobic). The lines represent all the Euclidean distances to be calculated for one residue. **B: Determination of the space point associated with a residue.** In the image, the spheres depict the different atoms of an amino acid, in this instance a glutamate residue. A cubic box is constructed as small as possible that contains the centers of every atom. Then, the geometrical center of the cubic box (yellow point) is considered as the space point for the residue.

On the other hand, the chemical nature of each residue corresponds to that of its R-chain. Specifically, Asp and Glu are designated as acidic, whilst Arg, His and Lys as basic. Polar residues include Ser, Thr, Asn and Gln. Finally, hydrophobic residues comprise Ala, Val, Ile, Leu, Met, Phe, Tyr and Trp. The rest of amino acids (Cys, Sec, Gly, Pro) are not included in the analysis.

Once the space point and chemical nature for each residue have been established, the calculation of QUILLO can proceed. The basic measurement involves the determination of Euclidean distances between residues of the same kind. That is, the minimum distance between the space point of residues of the same chemical nature. Figure 1A shows an illustration of this.

For every residue, the Euclidean distance with every other residue of the same type in the protein is calculated. Then, the inverse of each distance is squared, and the are results added up, rendering QUILLO. A formal definition can be seen in Equation 1.

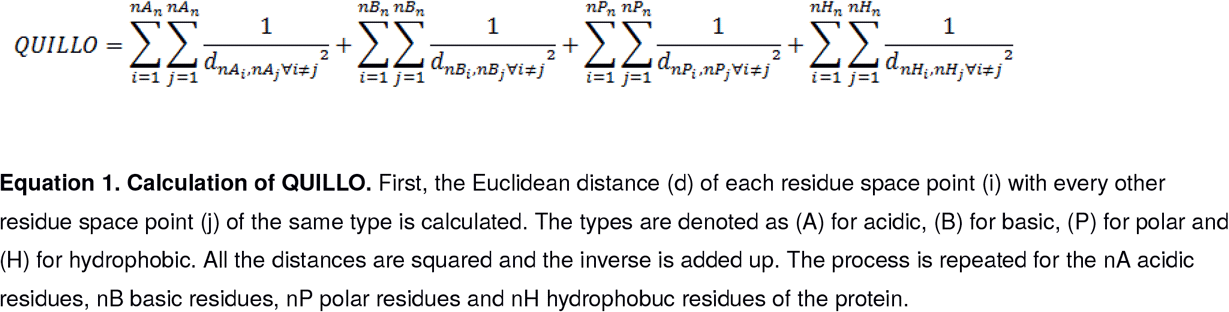

An alternative parameter (QUILLO2) is also presented in this work. Although being analogous to QUILLO, it does not include chemical nature specification. So, its definition --contained in Equation 2-- becomes much simpler. As discussed later, QUILLO2 offers less predictive power than QUILLO.

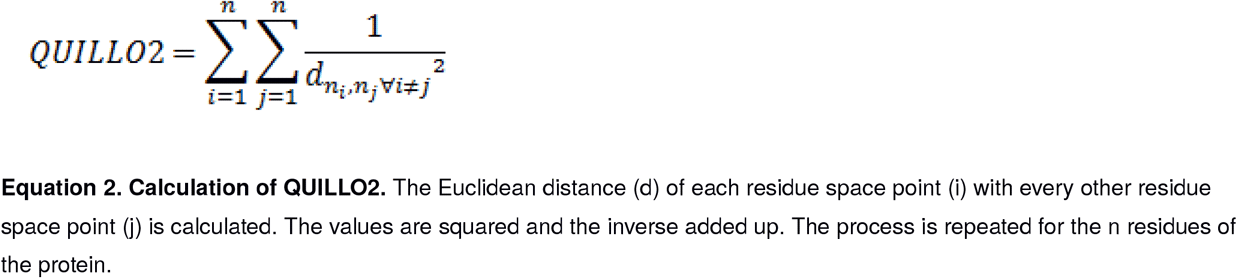

Both QUILLO and QUILLO2 constitute measurements of residue spatial clustering. By measuring the inverse of squared distances, their value becomes higher for increased amino acid clustering. In the calculations, Bio.PDB [38] [39] was used to extract the atom coordinates of each residue.

### Calculation of secondary structure composition

The DSSP database and related program mkdssp [17] [18] were employed as a dependency. In this manner, the percentage of residues in beta strand of each X-Ray PDB entry was obtained.

### Calculation of protein compactness

To evaluate protein compactness, the following strategy was followed. First, a sphere of minimum volume was generated for each protein that contained all its atoms. Then, the number of residues per sphere volume unit; i.e residue density in the sphere, was calculated. Since the spherical form is the most compact shape, the calculated residue density constitutes a direct measure of compactness for each entry.

### Calculation of surface area per residue

To compute surface areas of proteins, the software 3V [40] was used as a dependency. Probe radius was set to 1.5 Å. Subsequently, the data were normalized by the number of residues.

### Assessment of CASP12 protein predictions using the QUILLO proportion

Prediction structures and their corresponding rankings were obtained obtained from: http://predictioncenter.org/download_area/CASP12/.

For all the structures, the theoretical and real QUILLO value was calculated. Finally, the deviation from expected and real QUILLO was plotted against the structure’s ranking.

### Calculation of system temperature

System temperature refers here to the normal temperature in which the protein is found in nature. Calculating this value required making several approximations. Firstly, it was assumed that its value matched the species optimum growth temperature. In this way, all the proteins produced by a species would have the same system temperature. Secondly, that optimum growth temperature would coincide with the average optimum temperature of the species’ enzymes. The BRENDA database [19][20] provided the optimum temperatures of each species’ enzymes. These approximations, along with the use of the BRENDA database, enabled the calculation of system temperature for a total of 87,061 PDB entries.

### GO-term analysis of the QUILLO proportion

First, all X-Ray PDB entries were ordered according to their deviation from expected QUILLO values. Then, two groups were selected; a) those entries under percentile 10, and b) those above percentile 90. The GO-terms present in these two groups were extracted, and Fisher tests were carried out for each GO-term. In this way, a series of overrepresented GO-terms and underrepresented GO-terms were obtained for groups a) and b).

## Discussion

The present work shows a universal law of protein folding. This characteristic does not depend on secondary structure content, surface area or compactness. Instead, the QUILLO parameter depends just on the number of residues, following a polynomial regression. Along with the nature of the peptide bond, the presence of 20 standard amino acids or the organization in domains, this could be one of the few constant characteristics in proteins.

The reported proportion can be employed to assess protein structure predictions. To test this, data from the last CASP experiment was used. In effect, low ranks in the experiment corresponded to unexpected proportion values.

Also, the QUILLO proportion may open a new way of understanding polypeptides. Thermodynamically, the parameter could reflect high stable conformations. Biologically, it could be associated with protein dynamics, affecting in this way protein function. When comparing polypeptides with lower and higher QUILLO than expected, it is shown that catalytic activity is underrepresented in the first group and overrepresented in the second.

## Acknowledgements

I thank Fernando Govantes, Alejandro Cuetos, Juan Neftalí Morillo García and Francisco Javier García Moscoso for their technical support.

